# Parkinson’s disease-associated LRRK2 risk variant, G2385R, enhances Rab substrate phosphorylation and impairs neuronal integrity

**DOI:** 10.64898/2026.04.23.720426

**Authors:** An Phu Tran Nguyen, Roger Moser, Nicole Bryant, Lindsey A. Cunningham, Evan J. Worden, Darren J. Moore

## Abstract

Mutations in the *LRRK2* gene are the most frequent cause of familial Parkinson’s disease (PD) whereas common variants are associated with an increased risk for sporadic PD. *LRRK2* encodes a multi-domain protein displaying two functional enzymatic activities: GTPase and kinase. Familial *LRRK2* mutations have been linked to alterations in its GTPase and kinase activities, elevated substrate phosphorylation, as well as increased neurotoxicity in cell and animal models. In addition to familial mutations, several common *LRRK2* coding variants have been associated with PD risk in different ethnic populations. Little is known about how these coding variants modulate the risk of developing PD. In this study, we aim to evaluate whether a collection of *LRRK2* coding risk variants (A419V, N551K, R1398H, R1628P, M1646T, S1647T, G2385R) modify the biochemical properties of the LRRK2 protein and LRRK2-mediated neurotoxicity using cell-based assays. With the exception of G2385R, we find that coding risk variants have minimal impact on LRRK2 steady-state protein levels, GTP-binding, phosphorylation at Ser910, Ser935, and Ser1292, and subcellular localization in human cell lines and cultured primary neurons. G2385R reduces the steady-state levels of LRRK2 and exhibits reduced phosphorylation at Ser910/Ser935 and Ser1292, consistent with diminished kinase activity. Notably, however, *LRRK2* variants associated with increased PD risk (A419V, R1628P, M1646T, and G2385R) significantly elevate the levels of LRRK2-mediated Rab10 phosphorylation by two-fold in cells. In contrast, *LRRK2* variants associated with reduced PD risk (N551K, R1398H, and N551K-R1398H) do not alter normal LRRK2-mediated pRab10 levels. Intriguingly, we find that both PD-risk and PD-protective LRRK2 variants can respond normally to kinase activation by different lysosomal stressors (chloroquine, nigericin and monensin) to robustly induce pRab10 levels in cells. The PD risk variant, G2385R LRRK2, significantly inhibits neurite outgrowth in primary cortical neurons compared to wild-type LRRK2. Combining each coding risk variant with the familial G2019S mutation has minimal impact on the elevated kinase activity of G2019S LRRK2 or its capacity to inhibit neurite outgrowth. Our study indicates that LRRK2-dependent Rab phosphorylation represents a relevant readout of PD risk induced by *LRRK2* coding variants and demonstrates that the PD risk variant, G2385R LRRK2, creates a hyperactive kinase that can impair neuronal integrity at comparable levels to the effects of G2019S LRRK2.

## Introduction

Mutations in the *LRRK2* gene, which encodes leucine-rich repeat kinase 2, are the primary cause of late-onset, autosomal dominant Parkinson’s disease (PD) [1, 2]. At least seven familial mutations (N1437H, R1441C/G/H, Y1699C, G2019S, and I2020T) have been confirmed as pathogenic based upon their segregation with disease in families. These mutations are relatively rare and exhibit varying prevalence among different populations, suggesting ethnic-specific and geographical differences in *LRRK2* variant distribution [3-7]. *LRRK2* encodes a large protein (∼280 kDa) with multiple domains including armadillo repeats (ARM), ankyrin repeats (ANK), leucine-rich repeats (LRR), a short hinge-helix, a Ras-of-complex (Roc) GTPase domain followed by two tandem fold C-terminal-of-Roc (COR-A and COR-B) domains, a catalytic kinase domain, a WD40 domain, and an extended C-terminal αC-helix [8]. Familial mutations occur within the Roc GTPase (R1441C/G/H, N1437H), COR (Y1699C), or kinase (G2019S, I2020T) domains, implicating altered LRRK2 enzymatic activity in PD pathogenesis. Mutations in the kinase domain (G2019S, I2020T) directly increase LRRK2 kinase activity *in vitro*, whereas mutations in the Roc-COR domain (R1441C/G/H, Y1699C) impair GTP hydrolysis and/or increase GTP binding [9]. LRRK2 can autophosphorylate at Ser1292 located at the C-terminal end of the LRR domain and can phosphorylate substrate proteins in cells, including a subset of Rab GTPases (i.e. Rab8a, Rab10, Rab12) [10, 11]. Each of the seven familial LRRK2 mutations consistently increase Ser1292 autophosphorylation and Rab substrate phosphorylation at a specific site within the effector binding Switch II motif, indicating enhanced kinase activity [11]. Furthermore, some familial mutations (R1441C/G, Y1699C, I2020T) can decrease phosphorylation of LRRK2 at Ser910 and Ser935, reduce 14-3-3 protein binding, and enhance microtubule association [12]. Phosphorylation of LRRK2 at the biomarker sites, Ser910, Ser935, Ser955 and Ser973, serves as an indirect indicator of the conformation of the kinase domain and mediates the interaction with 14-3-3 proteins [13, 14]. Activation of LRRK2 involves its recruitment to cellular membranes by Rab GTPases via the N-terminal ARM domain [15]. Lysosomal stressors, such as monovalent cation ionophores and lysosomotropic agents, promote the relocalization of LRRK2 to lysosomal membranes where it increases Rab phosphorylation [16-19]. Elevated LRRK2 kinase activity can contribute to neuronal cell death, impaired neurite outgrowth, and disruption of Golgi complex morphology in primary neuronal models and certain animal models [20]. As PD-linked mutations consistently elevate LRRK2 kinase activity, selective LRRK2 kinase inhibitors are being tested in clinical trials as therapeutics for PD [21].

In addition to the familial *LRRK2* mutations, several common *LRRK2* coding variants are independently associated with modulating the lifetime risk of sporadic PD [22, 23] (**Table 1** and **Figure 1**). Among these variants, A419V, R1628P, S1647T, and G2385R, are frequently observed to increase PD risk in East Asian populations but are either absent or extremely rare in Caucasians [7, 23-26]. Another variant, M1646T, is associated with increased PD risk in Caucasians but not in Asian populations [23]. Interestingly, a common haplotype containing the nonsynonymous substitutions, N551K and R1398H, along with a synonymous substitution K1423K, is linked to reduced PD risk and is found across different populations [23]. Several of these variants have been associated with altered LRRK2 enzymatic activity *in vitro* and in cells [27-29]. However, the mechanism by which *LRRK2* coding risk variants influence susceptibility to sporadic PD remains unclear.

**Figure 1.**
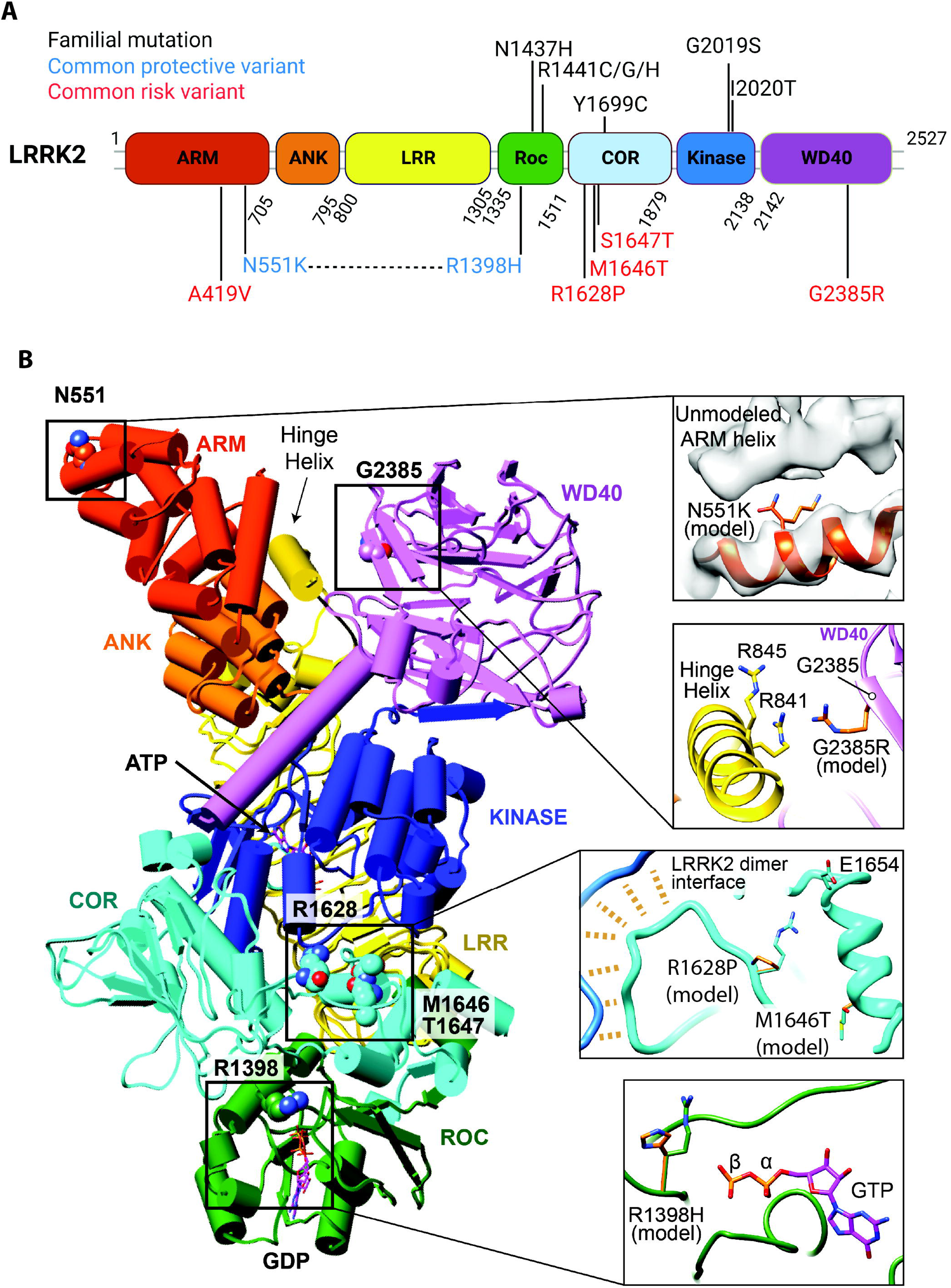
Domain architecture of human LRRK2 and impact of LRRK2 common coding variants on LRRK2 structure. (A) LRRK2 is shown as a multi-domain protein containing two key enzymatic domains: a Roc-GTPase domain and a kinase domain. LRRK2 common coding variants are labeled in blue (PD-protective variant) or red (PD-risk variant) and confirmed pathogenic mutations are labeled in black. (B) Structure of LRRK2 [558-2527] with depicted as pipes and planks with domains colored as in (A). Detailed views of LRRK2 variants at residues N551 (PDB:7LHW), R1398 (PDB:7LHW), R1628 (PDB:7LHT), M1646 (PDB:7LHT) and G2385 (PDB:7LHW) are shown in stick representation. Models of the mutated residues are colored orange. LRRK2 dimer contacts (PDB:7LHT) are indicated with the dashed orange lines.

**Table 1.**
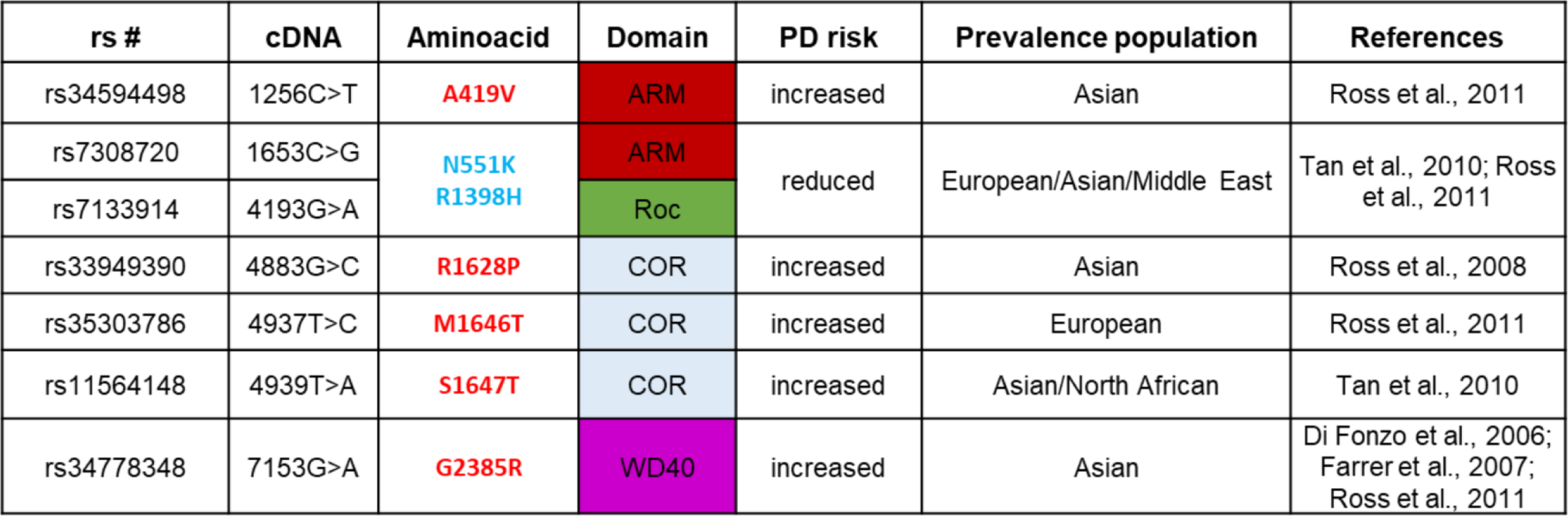
Common LRRK2 coding variants associated with PD risk. For each coding missense variant, the corresponding SNP rs number, nucleotide change within the cDNA, and domain location are shown. The direction of PD risk (increased or reduced) and the affected risk population are indicated.

Here, we investigate the impact of putative PD-risk variants (A419V, R1628P, M1646T, S1647T, and G2385R) and PD-protective variants (N551K, R1398H, and N551K-R1398H) on the biochemical properties of LRRK2 and its cellular activities with and without lysosomal stress. Additionally, we evaluate whether PD risk variants of LRRK2 can inhibit neurite outgrowth, and whether PD protective variants can mitigate neurite outgrowth inhibition induced by familial G2019S LRRK2 in primary cortical neurons. Together, this study provides a comprehensive functional characterization of *LRRK2* coding risk variants associated with PD in mammalian cell lines and neurons.

## Results

### Predicted impact of coding risk variants on LRRK2 structure

To explore the mechanisms by which *LRRK2* coding variants may influence PD risk, we first analyzed their predicted impact on LRRK2 structure using high-resolution structures of inactive full-length LRRK2 (Protein Data Bank (PDB) 7LHW and 7LHT [8]). The coding risk variants selected for this study are located across different domains of LRRK2, including the ARM domain (A419V, N551K), Roc-GTPase domain (R1398H), COR-A domain (R1628P, M1646T, S1647T), and WD40 domain (G2385R) (**Figure 1A**). The A419V variant is absent from the currently available structure. We find that the N551K variant may disrupt contacts within the unmodeled N-terminal ARM domain. The R1398H variant likely affects GTPase activity, possibly through interactions with gamma phosphate of GTP. The COR-A variants, R1628P and M1646T, might disrupt the structure of a loop that mediates interactions within the LRRK2 dimer. The Ser1647 residue has already been replaced by Threonine in the present model, so the impact of the S1647T variant cannot be analyzed. Interestingly, the G2385R variant likely causes charge repulsion with residues R841 and R845 located in the hinge-helix, which may destabilize the WD40 domain (**Figure 1B**).

### Impact of coding risk variants on LRRK2 steady-state levels, GTP-binding, and phosphorylation at Ser910 and Ser935 in cells

To evaluate the functional impact of coding risk variants on the biochemical properties of LRRK2, we generated full-length FLAG-tagged human LRRK2 expression constructs containing individual PD risk variants (A419V, R1628P, M1646T, S1647T, and G2385R) or protective variants (N551K, R1398H, N551K-R1398H) (**Figure 1** and **Table 1**). We first analyzed the effects of each risk variant on the steady-state protein levels of LRRK2 transiently expressed in SH-SY5Y cells. We find no significant changes in the LRRK2 steady-state levels of risk variants compared to wild-type (WT) LRRK2, although G2385R is consistently reduced (**Figure 2A, B**). We next assessed the impact of risk variants on LRRK2 GTPase activity by monitoring its GTP-binding capacity. Pull-down assays using GTP-agarose on cell extracts expressing LRRK2 variants reveal that risk variants do not exhibit significant differences in the steady-state levels of GTP-binding compared to WT LRRK2 (**Figure 2C, D**), although N551K and R1398H are modestly increased. A known GTP-binding-deficient mutation (T1348N) serves as a negative control for this assay [30]. Notably, the protective variants, R1398H and N551K-R1398H, have been reported to impair GTP-binding or increase GTP hydrolysis [28, 31], although we are not able to replicate these data. The constitutive phosphorylation status of LRRK2 at the biomarker residues Ser910 and Ser935 (pSer910-LRRK2 and pSer935-LRRK2) was next evaluated. Western blot analysis of detergent-soluble extracts from SH-SY5Y cells transiently expressing each LRRK2 risk variant, indicates that G2385R significantly reduces Ser910 and Ser935 LRRK2 phosphorylation, with a modest reduction by A419V, compared to WT LRRK2 and other risk variants (**Figure 2E-G**). The familial R1441C mutation, and kinase-inactive T1348N and D1994A mutations, also exhibit reduced Ser910/Ser935 phosphorylation as reported [13, 30], and serve to calibrate these assays. Of note, G2385R and R1441C induce comparable reductions in LRRK2 phosphorylation (**Figure 2E-G**).

**Figure 2.**
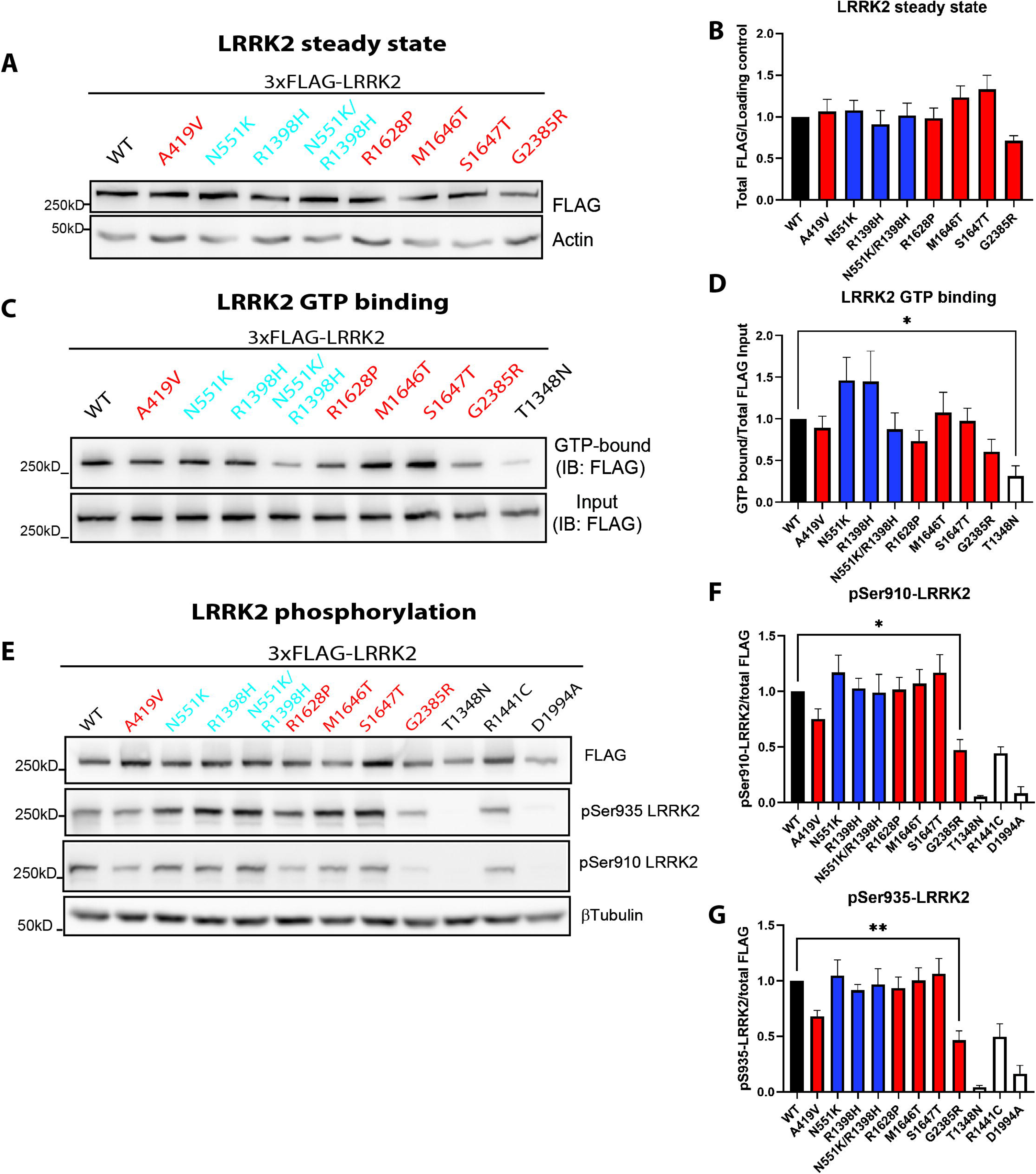
Impact of coding risk variants on LRRK2 steady-state levels, GTP-binding, cellular and Rab10 phosphorylation in cells. (A) Western blot analysis of steady-state levels of FLAG-tagged LRRK2 risk variants transiently expressed in SH-SY5Y cells. (B) Densitometric analysis of LRRK2 levels normalized to actin expressed as a proportion of WT LRRK2. Bars represent mean ± SEM levels of FLAG-LRRK2 (*n* = 10 independent transfections). (C) Western blot analysis of GTP-binding capacity of FLAG-tagged LRRK2 risk variants or synthetic mutations relative to input levels in SH-SY5Y cells. (D) Densitometric analysis of GTP-bound LRRK2 normalized to FLAG-LRRK2 input levels expressed as a proportion of WT LRRK2. Bars represent mean ± SEM of GTP-bound LRRK2 (*n* = 7 independent transfections). (E) Western blot analysis of constitutive phosphorylation of FLAG-tagged LRRK2 risk variants or familial/synthetic mutations at Ser910 and Ser935 in SH-SY5Y cells. Densitometric analysis of (F) pSer910-LRRK2 or (G) pSer935-LRRK2 levels normalized to total FLAG-LRRK2 expressed as a proportion of WT LRRK2. Bars represent mean ± SEM of LRRK2 phosphorylation (*n* = 10 independent transfections). \**P*<0.05 or \*\**P*<0.01 compared with WT LRRK2 by one-way ANOVA with matching rows and Dunnett’s multiple comparison test.

### PD risk variants enhance LRRK2-mediated Rab10 phosphorylation but not LRRK2 autophosphorylation

The kinase activity of LRRK2 coding risk variants was examined by monitoring the phosphorylation levels of two *bona fide* LRRK2 substrates: LRRK2 itself via autophosphorylation at Ser1292 (pSer1292-LRRK2) and Rab10 at Thr73 (pThr73-Rab10) [10, 11]. Western blot analysis using phospho-specific antibodies reveals no significant differences in pSer1292-LRRK2 levels between coding risk variants and WT LRRK2 (**Figure 3A, B**), although there is a trend for G2385R to be reduced, whereas familial R1441C increases and kinase-inactive T1348N or D1994A mutations reduce pSer1292 levels, as expected. To measure pThr73-Rab10, we co-transfected GFP-tagged Rab10 with each FLAG-tagged LRRK2 variant in SH-SY5Y cells and performed Western blot analysis with a phospho-specific antibody for pThr73-Rab10 [11, 27, 29]. Intriguingly, we find that PD risk variants (A419V, R1628P, M1646T, and G2385R) increased pT73-Rab10 levels by ∼2-fold compared to WT LRRK2 (**Figure 3A, C**), with a more modest increase for the S1647T variant. PD protective variants (N551K, R1398H, N551K-R1398H) have no significant effect on pThr73-Rab10 levels compared to WT LRRK2, although the N551K variant alone or combined modestly reduces pRab10 (**Figure 3A, C**). As expected, the familial R1441C mutation robustly increases LRRK2-mediated Rab10 phosphorylation by ∼3-fold, while kinase-inactive T1348N and D1994A mutations have negligible effects, thereby confirming the specificity of these assays (**Figure 3A, C**). Endogenous pThr73-Rab10 in SH-SY5Y cells is not easily detected due to low levels of Rab10, hence the use of exogenous GFP-Rab10 in these cells. Collectively, our data demonstrate that PD risk variants promote LRRK2 kinase activity towards its cellular substrates, as revealed by pThr73-Rab10 levels.

**Figure 3.**
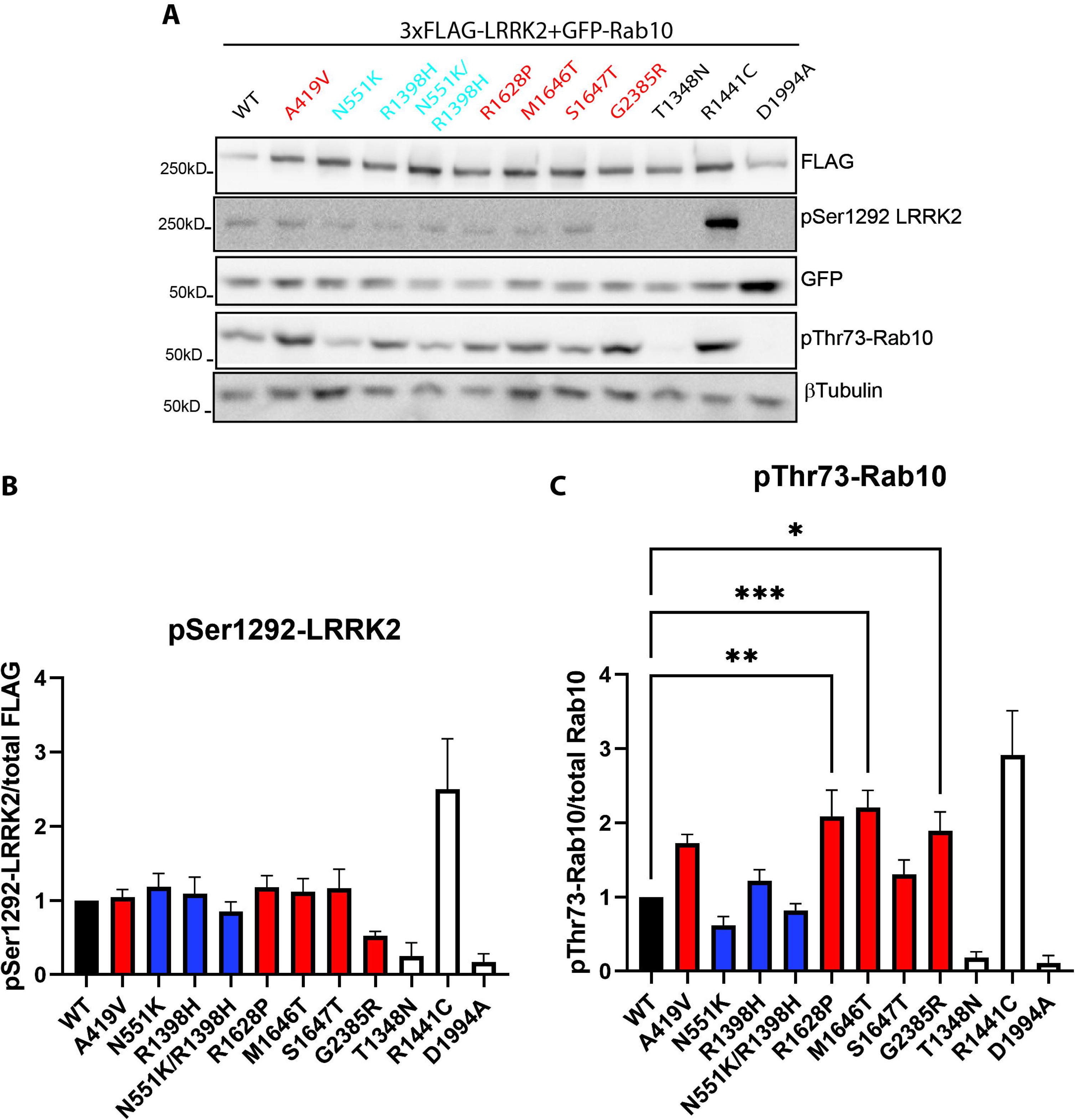
Impact of LRRK2 coding risk variants on Ser1292 autophosphorylation and Rab10 phosphorylation in cells. (A) Western blot analysis of LRRK2 autophosphorylation at Ser1292 and phosphorylation of GFP-Rab10 at Thr73 induced by FLAG-tagged LRRK2 risk variants or familial/synthetic mutations in SH-SY5Y cells. (B) Densitometric analysis of pSer1292-LRRK2 levels normalized to total FLAG-LRRK2 expressed as a proportion of WT LRRK2. Bars represent mean ± SEM levels of pSer1292-LRRK2 (*n* = 8 independent transfections). *P*>0.05 compared with WT LRRK2 by one-way ANOVA with matching rows and Dunnett’s multiple comparison test. (C) Densitometric analysis of pThr73-Rab10 levels normalized to total GFP-Rab10 expressed as a proportion of WT LRRK2. Bars represent mean ± SEM levels of pThr73-Rab10 (*n* = 8 independent transfections). \**P*<0.05, \*\**P*<0.01 or \*\*\**P*<0.005 compared with WT LRRK2 by one-way ANOVA with matching rows and Dunnett’s multiple comparison test.

### Normal cytoplasmic localization of LRRK2 coding risk variants in cell lines and neurons

LRRK2 normally adopts a diffuse cytoplasmic localization in cells, but abnormal LRRK2 activity can alter its subcellular localization [13]. Familial LRRK2 mutants (R1441C/G/H, Y1699C, I2020T) and kinase inhibitor-treated LRRK2 exhibit reduced pSer910 and pSer935 levels and preferentially form filamentous skein-like structures decorating microtubules when overexpressed in cells [12, 32, 33]. To assess the impact of coding risk variants on LRRK2 localization, we examined HEK-293T cells and rat primary cortical neurons transiently expressing LRRK2 variants by immunocytochemistry and confocal microscopy. We find that both PD-risk and PD-protective LRRK2 variants exhibit equivalent diffuse cytoplasmic localization in HEK-293T cells compared to WT LRRK2 and the familial G2019S mutant (**Figure S1**). Co-labeling of LRRK2 variants with endogenous pThr73-Rab10 reveals consistent overlap in HEK-293T cells, confirming LRRK2 kinase activation (**Figure S1**). In rat cortical neurons, we again observe an equivalent diffuse cytoplasmic localization of PD-risk and PD-protective LRRK2 variants compared to WT LRRK2 without obvious cytoplasmic inclusions or enhanced LRRK2-positive filament formation, including for familial Y1699C LRRK2 (**Figure S2**). In cortical neurons, we find negligible levels of endogenous pThr73-Rab10 labeling, even in cells expressing kinase-enhancing familial LRRK2 mutations or PD risk variants (**Figure S2**), most likely indicating a low abundance of Rab10 in these neuronal subtypes. Although most familial *LRRK2* mutations promote the formation of microtubule-bound LRRK2 filaments in cell lines when overexpressed, this feature has not been extensively evaluated in neuronal cell types. To further explore the capacity of LRRK2 risk variants to form filaments in neurons, we treated rat primary neurons transiently expressing LRRK2 variants with the LRRK2 kinase inhibitor MLi-2 (100 nM) for 3 hours (**Figure S3**). We find that neither the familial Y1699C LRRK2 mutant, nor the PD-risk variants A419V and G2385R, are able to form LRRK2-positive filamentous structures following MLi-2 treatment in cortical neurons (**Figure S3**). MLi-2 treatment does efficiently reduce exogenous and endogenous pSer935-LRRK2 levels in cortical neurons, thereby confirming LRRK2 kinase inhibition (**Figure S3**). These data indicate that LRRK2 coding risk variants are not sufficient to induce cytoplasmic inclusion formation in different cells.

### Coding risk variants induce normal LRRK2 activation in response to lysosomal stress

Treatment with lysosomal stressors, such as monovalent cation ionophores (nigericin and monensin) and lysosomotropic agents (chloroquine and L-leucyl-L-leucine methyl ester [LLOMe]), has been shown to stimulate LRRK2 kinase activity and elevate Rab10 substrate phosphorylation in cultured cells [16-19]. Given that PD-risk and PD-protective variants of LRRK2 induce different levels of Rab10 phosphorylation under normal conditions (**Figure 3A, C**), we sought to determine whether they respond differently to lysosomal stressors. We transiently co-expressed LRRK2 coding variants and GFP-Rab10 in HEK-293T cells for 48 hours, followed by treatment with lysosomal stress agents. We find that treatment with nigericin (2 μM) for 2 hours significantly increases pThr73-Rab10 levels up to 3-fold in cells expressing WT LRRK2 (**Figure 4A**). Treatment with chloroquine (50 µM) or monensin (10 μM) significantly increases Rab10 phosphorylation up to 2-fold in cells expressing WT LRRK2 (**Figures 4B** and **S4A**). The levels of pThr73-Rab10 similarly increase in cells expressing PD-risk or PD-protective variants of LRRK2 following treatment with nigericin, chloroquine, or monensin (**Figures 4** and **S4A**). For treatment with LLOMe, we limited treatment time to 1 hour to avoid excessive cell detachment. We find that LLOMe (1 mM) increases Rab10 phosphorylation by only ∼1.2-fold in cells expressing WT LRRK2, whereas coding risk variants produce a similar small increase or no effect on pThr73-Rab10 levels (**Figure S4B**). Interestingly, while lysosomal stressors elevate LRRK2-mediated Rab10 phosphorylation, they do not significantly alter the levels of pSer1292-LRRK2 or pSer935-LRRK2 in cells (**Figures 4** and **S4**). As controls in these assays, kinase-inactive D1994A LRRK2, and to a lesser extent familial Y1699C LRRK2, do not increase pThr73-Rab10 levels in response to lysosomal stress, whereas familial G2019S LRRK2 behaves similarly to LRRK2 coding risk variants and WT (**Figures 4** and **S4**). Collectively, our data confirm the efficacy of lysosomal stressors in inducing LRRK2-dependent Rab10 phosphorylation in cells, and demonstrate that both PD-risk and PD-protective LRRK2 variants can respond normally to lysosomal stress by enhancing Rab10 phosphorylation.

**Figure 4.**
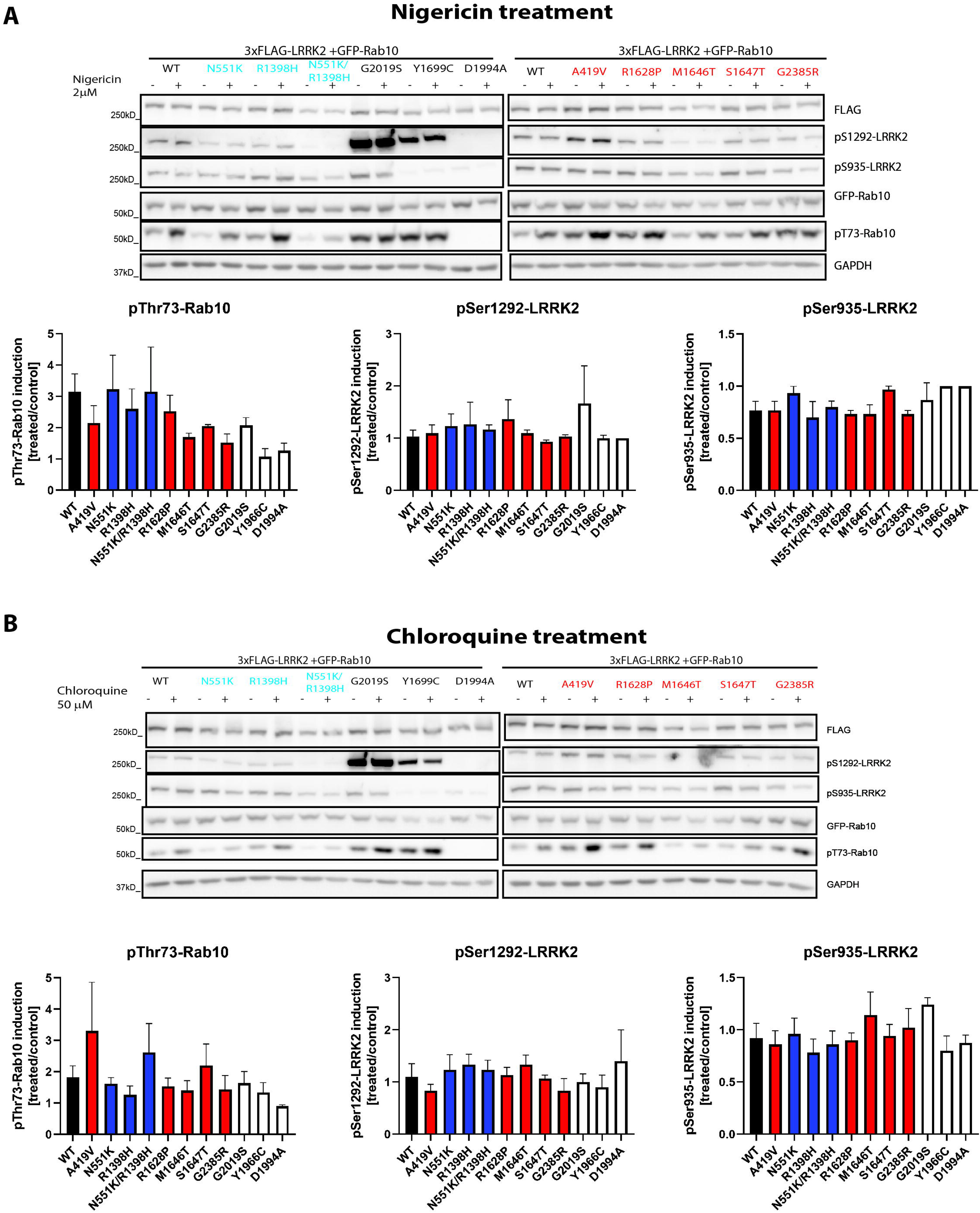
PD-risk and PD-protective LRRK2 coding variants support increased Rab10 phosphorylation induced by lysosomal stressors. Western blot analysis of GFP-Rab10 phosphorylation at Thr73 and LRRK2 phosphorylation at Ser935 and Ser1292 induced by FLAG-tagged LRRK2 risk variants or familial/synthetic mutations in SH-SY5Y cells treated with the lysosomal stressors (A) 2 µM nigericin or (B) 50 µM chloroquine relative to PBS (control) for 2 h. (A-B) Bars represent mean ± SEM fold induction of pThr73-Rab10, pSer1292-LRRK2, or pSer935-LRRK2 for each LRRK2 risk variant by each lysosomal agent relative to control treatment (*n* = 3-5 independent transfections). *P*>0.05 compared with WT LRRK2 by multiple Student’s *t*-test.

### Limited impact of coding risk variants on the kinase activity of PD-linked G2019S LRRK2 in cells

To further evaluate potential mechanisms of PD risk by LRRK2 coding variants, we sought to determine if they can modify the pathogenic effects of the familial G2019S LRRK2 mutation. To do this, we combined each LRRK2 coding risk variant with the G2019S mutation. We first assessed the impact of coding risk variants on G2019S LRRK2 steady-state levels, phosphorylation (Ser910 and Ser935), and kinase activity (pS1292-LRRK2 and pThr73-Rab10) in SH-SY5Y cells using methodologies as described above (**Figures 2, 3**). We find that coding risk variants do not influence the steady-state levels of G2019S LRRK2 protein in SH-SY5Y cells (**Figure 5A, B**). There are no significant differences in the levels of pSer910-LRRK2, pSer935-LRRK2, pSer1292-LRRK2, or pThr73-Rab10 in cells expressing combined LRRK2 coding variants/G2019S, compared to G2019S LRRK2 alone (**Figure 5A, C-F**). However, the PD risk variants (R1628P, M1646T, S1647T, G2385R) combined with G2019S tend to modestly increase pThr73-Rab10 levels compared to G2019S alone (**Figure 5F**). We also note that the G2385R variant can consistently reduce pS910/pS935 levels of G2019S LRRK2 (**Figure 5C, D**), similar to its impact on WT LRRK2 (**Figure 2F, G**). T1348N and D1994A LRRK2 serve as kinase-inactive controls in these assays. Collectively, our data demonstrate that LRRK2 coding risk variants are not able to robustly alter the biochemical properties or kinase activity of the familial G2019S LRRK2 mutant in cells. Importantly, PD-protective variants (N551K, R1398H or N551K-R1398H) are not able to lower the elevated kinase activity of G2019S LRRK2.

**Figure 5.**
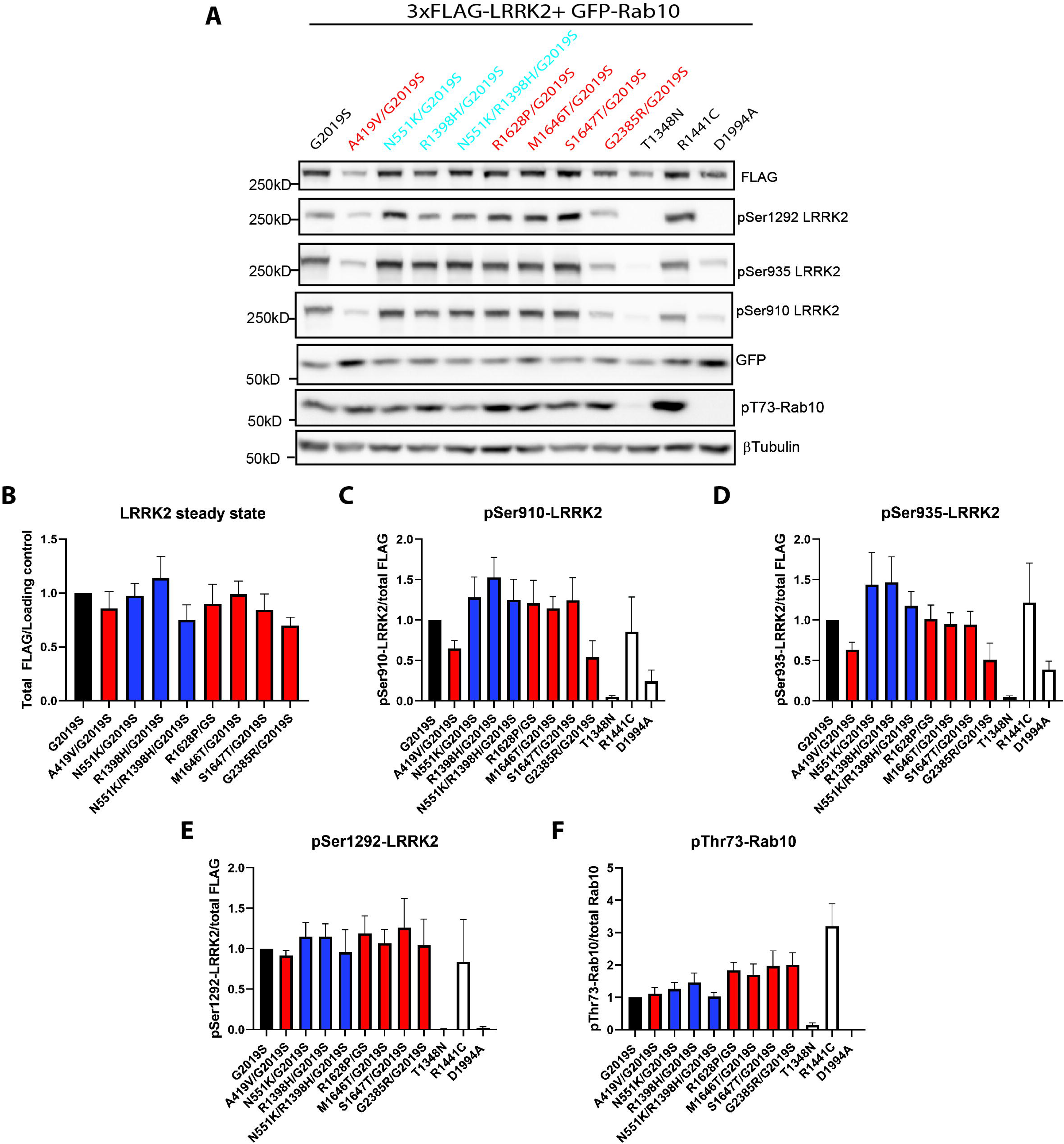
LRRK2 coding risk variants have minimal impact on the steady-state protein levels, cellular phosphorylation, and kinase activity of G2019S LRRK2. (A) Western blot analysis of LRRK2 steady-state levels, LRRK2 phosphorylation (at Ser910, Ser935, or Ser1292), and GFP-Rab10 phosphorylation at Thr73 induced by FLAG-tagged LRRK2 risk variants combined with the G2019S mutation in SH-SY5Y cells. (B) Densitometric analysis of FLAG-LRRK2 variant levels normalized to β-tubulin expressed as a proportion of WT LRRK2. Bars represent mean ± SEM (*n* = 5 independent transfections). (C-E) Densitometric analysis of LRRK2 phosphorylation (Ser910, Ser935, or Ser1292) normalized to total FLAG-LRRK2 expressed as a proportion of WT LRRK2. Bars represent mean ± SEM (*n* = 4 independent transfections). (F) Densitometric analysis of pThr73-Rab10 levels normalized to total GFP-Rab10 expressed as a proportion of WT LRRK2. Bars represent mean ± SEM (*n* = 4 independent transfections), *P*>0.05 compared with WT LRRK2 by one-way ANOVA with matching rows and Dunnett’s multiple comparison test.

### PD-risk variant G2385R impairs neurite outgrowth, whereas PD-protective variants fail to attenuate the neuronal effects of G2019S LRRK2

We next evaluated the effects of LRRK2 coding risk variants on neurite outgrowth in rat primary cortical neurons, as a surrogate readout of impaired neuronal integrity or neuronal toxicity. Prior studies have shown that G2019S LRRK2 robustly impairs neurite outgrowth in primary neurons, compared to WT LRRK2 which has negligible effects [30, 34]. In this assay, we measured the length of the longest neurite (i.e. the axon) in cortical neurons that transiently co-expressed each FLAG-tagged LRRK2 variant and GFP (FLAG+/GFP+ neurons), where GFP serves as a useful marker to identify neuronal processes. We find that neurons expressing WT LRRK2 exhibit comparable neurite length compared to control neurons (empty vector) (**Figure 6A, B**), while in similar assays G2019S LRRK2 significantly reduces neurite length by ∼40% compared to control neurons (**Figure 6C, D**), as previously reported [30, 34]. Interestingly, among PD risk variants, only the G2385R LRRK2 variant significantly reduces neurite length by ∼50% compared to WT LRRK2 or control neurons, suggesting that this variant alone induces neuronal toxicity (**Figure 6A, B**). The PD-risk variants, A419V, R1628P, M1646T, and S1647T, modestly reduce neurite length compared to WT LRRK2 (**Figure 6A, B**), albeit this was not significant in these assays. Somewhat unexpectedly, two PD-protective variants (N551K and N551K-R1398H) also significantly decrease neurite length compared to WT LRRK2 but to a lesser extent than G2385R (**Figure 6A, B**), whereas R1398H alone behaves similarly to WT LRRK2. We next combined each coding risk variant with G2019S LRRK2 to explore how they might alter its neurite outgrowth effects, and particularly whether protective variants can attenuate its pathogenic effects. Notably, however, we do not detect significant differences in neurite length in neurons expressing N551K/G2019S, R1398H/G2019S, or N551K-R1398H/G2019S LRRK2 compared to G2019S LRRK2 alone, with all variants similarly inducing a ∼40% reduction in neurite length compared to control neurons (**Figure 6C, D**). PD risk variants do not further exacerbate the neurite outgrowth inhibition induced by G2019S LRRK2, including G2385R, indicating their effects are neither additive nor synergistic (**Figure 6C, D**). These data indicate that PD-protective variants fail to rescue impaired neurite outgrowth induced by the familial G2019S LRRK2 mutant, whereas the G2385R risk variant alone impairs neuronal integrity.

**Figure 6.**
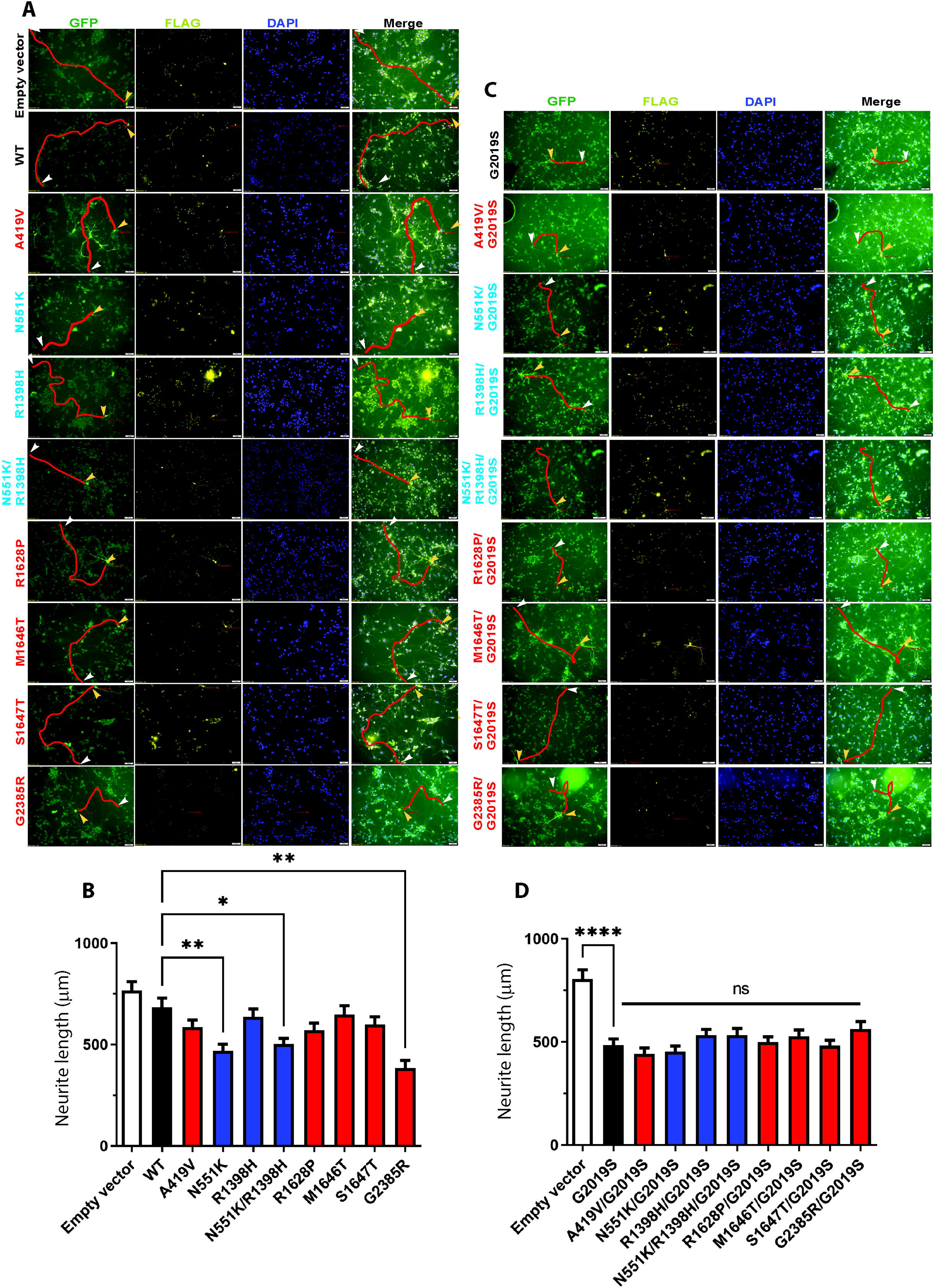
Impact of LRRK2 coding risk variants on neurite outgrowth inhibition induced by G2019S LRRK2 in primary cortical neurons. (A, C) Rat primary cortical neurons co-transfected at DIV 3 with each FLAG-tagged human LRRK2 variant (or empty control vector) and GFP plasmids at a DNA molar ratio of 10:1. Cultures were fixed at DIV 6 and labeled for immunofluorescence with anti-FLAG antibody. Shown are representative fluorescent micrographs revealing the co-labeling of cortical neurons with GFP (green), LRRK2 variants (FLAG, yellow) and nuclei (DAPI, blue). Measured GFP-positive neurites are traced in red and arrowheads indicate the start (yellow) and end (white) of the traced neurite. (B, D) Analysis of the length of GFP-positive neurites from FLAG-LRRK2-positive cortical neurons. (B) Bars represent mean ± SEM axon length (in µm) for each LRRK2 risk variant (*n* = 50-200 cells/condition from 2 independent cultures). \**P*<0.05 or \*\**P*<0.01 compared with cells expressing WT LRRK2 as indicated by one-way ANOVA with Dunnett’s multiple comparison test. (D) Bars represent mean ± SEM axon length for each LRRK2 risk/G2019S double mutant (*n* = 50-300 cells/condition from 3 independent cultures). \*\*\*\**P*<0.0001 compared with empty vector as indicated by one-way ANOVA with Dunnett’s multiple comparison test. Non-significant (*ns*, *P*>0.05) compared to cells expressing G2019S LRRK2 alone as indicated by one-way ANOVA.

## Discussion

The consistent elevation of LRRK2 kinase activity by PD-linked familial mutations supports the development of pharmacological LRRK2 kinase inhibitors as a potential therapeutic approach for *LRRK2*-linked PD. In addition to familial mutations, many common *LRRK2* coding variants are associated with PD risk. As LRRK2 kinase inhibitors are being evaluated in clinical trials, it is crucial to determine how common *LRRK2* coding variants modulate PD susceptibility, thus supporting precision medicine for this disease-modification approach. In this study, we characterize a series of common LRRK2 coding variants previously associated with increased risk (A419V, R1628P, M1646T, S1647T, and G2385R) or reduced risk (N551K, R1398H, N551K-R1398H) of developing sporadic PD. We find that the most common PD risk variant in East Asian populations, G2385R, reduces LRRK2 phosphorylation at Ser910 and Ser935, and autophosphorylation at Ser1292 in cells, apparently most consistent with a kinase-inactivating effect. In general, we find that LRRK2 variants associated with increased PD risk (A419V, R1628P, M1646T, S1647T, and G2385R) commonly increase Rab10 substrate phosphorylation up to 2-fold in cells. In contrast, there was no significant impact on pThr73-Rab10 levels induced by PD-protective variants (N551K, R1398H, and N551K-R1398H) in cells. Treatment with different lysosomal stressors consistently increased pRab10 levels in cells overexpressing either PD-risk or PD-protective LRRK2 variants, indicating that these coding variants retain the capacity for activation by lysosomal stress. Among PD risk variants, only G2385R LRRK2 expression significantly impairs neurite outgrowth in cortical neurons to a similar extent as the pathogenic G2019S mutation. LRRK2 coding risk variants are not able to modify the steady-state protein levels, phosphorylation, or kinase activity of G2019S LRRK2, or the inhibition of neurite outgrowth induced by G2019S LRRK2 in cortical neurons. Collectively, our study reveals that PD-risk variants tend to have subtle effects yet lead to a modest increase in LRRK2 kinase activity, suggesting that PD subjects harboring these variants (particularly G2385R carriers) might benefit from treatment with LRRK2 kinase inhibitors.

We note that the increase in pThr73-Rab10 levels induced by LRRK2 risk variants are moderate (up to 2-fold) compared to the effects of most familial LRRK2 mutations (2-4-fold), consistent with previous reports [27, 29]. Intriguingly, LRRK2 risk variants do not promote Ser1292 autophosphorylation, probably due to low stoichiometry phosphorylation at this site [15]. A recent report also found increases in Rab phosphorylation, but not pSer1292-LRRK2 levels, mediated by A419V, R1628P, and G2385R LRRK2 [29]. Supporting the relevance of our findings, a recent cohort study of Malaysian PD patients carrying the G2385R variant, revealed modestly increased pRab10 levels (∼1.2-fold) and reduced pSer935-LRRK2 levels (∼0.9-fold) in peripheral blood monocytes [35]. Our data replicates and extends these observations to additional LRRK2 risk variants and functional assays. In the case of the G2385R variant, we find reduced levels of Ser1292 phosphorylation yet increased Rab10 phosphorylation, suggesting that these two kinase-dependent events are uncoupled and implying that pSer1292-LRRK2 may not provide a reliable marker for kinase activation status. G2385R is the only risk variant we studied located in the WD40 domain, suggesting that this domain plays an important structural role in regulating Ser1292 autophosphorylation and/or kinase activation. We find no significant changes in Ser1292-LRRK2 autophosphorylation or Rab10 phosphorylation induced by PD-protective LRRK2 variants, consistent with previous reports [27-29, 36], although we find a trend towards reduced pThr73-Rab10 levels with the N551K and N551K-R1398H variants. It is not known which LRRK2 variant (N551K, R1398H, or K1423K), or combination of variants, within this haplotype region is associated with reduced PD risk, and our data might support a more prominent role for N551K over R1398H. While LRRK2 familial and coding risk variants consistently increase pThr73-Rab10 levels in cell lines, we find that endogenous pThr73-Rab10 levels are not abundant in primary cortical neurons (**Figure S2**), perhaps suggesting that other LRRK2 substrates are involved following LRRK2 activation in these neurons. Future studies should evaluate the impact of LRRK2 coding risk variants on the phosphorylation of different Rab substrates in neurons versus glial cells. Indeed, it has been suggested that some LRRK2 PD variants selectively enhance Rab10 phosphorylation without impacting Rab12 phosphorylation [29]. The impact of coding risk variants on other putative LRRK2 substrates, i.e. ArfGAP1, Endophilin A, MARK1, in cells and neurons, remains to be investigated. Collectively, our data reconcile the pathogenicity prediction of LRRK2 coding risk variants based on their REVEL score [22] with their impact on inducing Rab10 phosphorylation levels. We propose that Rab phosphorylation, but not LRRK2 autophosphorylation at Ser1292, is a more reliable indicator of LRRK2 kinase activity in cells and serves as the most useful measure of PD risk.

Although the G2385R variant was previously found to reduce LRRK2 kinase activity *in vitro* [37-39], our results reveal that it promotes Rab10 phosphorylation in cells, consistent with recent reports [11, 27, 29, 35, 40, 41]. Despite increased substrate phosphorylation, G2385R LRRK2 displays reduced Ser1292 phosphorylation. The G2385R variant is located at the dimer interface of WD40, a domain critically required for LRRK2 kinase activity and LRRK2-induced neuronal damage [42]. The G2385R variant disrupts WD40 dimers and the WD40-hinge helix interaction [8, 40, 41]. These conformational changes may destabilize the inactive state of LRRK2, resulting in more access for Rab proteins to be phosphorylated by LRRK2 [15]. Accordingly, we found that G2385R LRRK2 displays a significant reduction in phosphorylation at Ser910 and Ser935, suggesting an active (closed) conformation of the kinase domain, similar to most familial LRRK2 mutations [14]. Interestingly, overexpressing G2385R LRRK2 in cells or neurons does not lead to the formation of filamentous structures following treatment with LRRK2 kinase inhibitors, as previously reported [29], suggesting that an intact WD40 dimerization interface is required for the formation of LRRK2 filaments [12]. Our results demonstrate that the G2385R variant, despite reducing pSer910/935-LRRK2 levels, does not lead to the formation of microtubule-bound LRRK2 filaments in HEK-293T cells and rat cortical neurons (**Figures S1**, **S2**, and **S3**). However, the familial Y1699C mutation, which also exhibits reduced pSer910/935-LRRK2 levels, does not enhance filament formation in neurons in these assays. Additionally, treatment with a LRRK2 kinase inhibitor does not promote filament formation in neurons expressing LRRK2 risk variants (**Figure S3**). The pathogenic mechanism of LRRK2 variants in this neuronal cell type remains to be explored.

Among the PD risk variants, we found that only G2385R LRRK2 induces neurite outgrowth inhibition in primary cortical neurons, to a level comparable to the G2019S mutation. However, the G2385R variant did not have additive or synergistic effects on neurite outgrowth inhibition when combined with the G2019S mutation, implying that G2019S has a dominant effect on this neuronal phenotype and on LRRK2 substrate phosphorylation. Overexpression of G2385R LRRK2 induces centrosomal cohesion deficits in HEK-293T cells in a kinase-dependent manner [43]. In neurons, the G2385R variant altered the strength and quality of interactions with synaptic vesicles [44, 45]. Additionally, overexpression of G2385R LRRK2 in cells increases mitochondrial ROS levels and sensitizes neurons to oxidative stress-induced neurite injury [46]. Although we find no differences in Rab10 phosphorylation induced by lysosomal stressors in HEK-293T cells expressing G2385R LRRK2 compared to WT LRRK2, the effects of lysosomal damage in neurons expressing G2385R LRRK2 have not been explored. Similarly, a previous study has shown that G2385R LRRK2 also responds normally by increasing Rab10 phosphorylation in response to Rab29 overexpression [29]. The impact of exposing neurons expressing G2385R LRRK2 to protein aggregation (i.e., α-synuclein pre-formed fibrils) or environmental toxins linked to PD (i.e., rotenone, trichloroethylene) could be explored to determine whether G2385R LRRK2 carriers would be sensitized to these PD risk factors. Future studies should investigate the mechanism by which G2385R LRRK2 induces neuronal toxicity and neurodegeneration *in vivo*, especially the role of different LRRK2 enzymatic activities and particularly Rab substrate phosphorylation.

We find that the presumed protective variants (N551K, R1398H, or N551K-R1398H) do not mitigate the neurite outgrowth inhibition induced by G2019S LRRK2 in primary cortical neurons, again supporting the dominant effect of the G2019S mutation. When expressed alone, we find that N551K and N551K-R1398H LRRK2 significantly reduced neurite outgrowth compared to WT LRRK2, but to a lesser extent than G2385R or G2019S LRRK2. This was unexpected given that both of these LRRK2 variants also showed a trend towards reduced Rab10 phosphorylation. We speculate that these protective variants might exhibit intrinsic toxicity when overexpressed in neurons that might be absent at endogenous expression levels. The R1398H variant, located in the switch II motif of the Roc domain, has previously been reported to reduce GTP-binding, increase GTP hydrolysis, and enhance Roc-COR dimerization, opposite to the effects of familial mutations in the Roc-COR domain (i.e. R1441C/G/H and Y1699C) [28, 31]. Our study fails to replicate the impact of the R1398H variant on GTP-binding capacity and kinase activity. Our data reveal that N551K, R1398H, and N551K-R1398H LRRK2 do not significantly reduce Rab10 phosphorylation (despite a trend towards reduced pRab10 levels), autophosphorylation at Ser1292, or LRRK2 phosphorylation (Ser910, Ser935) compared to WT LRRK2, indicating that these protective variants do not obviously alter LRRK2 kinase activity in cells, largely consistent with previous reports [27-29]. A recent report found that N551K-R1398H LRRK2 significantly reduced Rab phosphorylation levels in cells compared to WT LRRK2, and our data lends some support to this observation. Notably, the total levels of N551K-R1398H LRRK2 were reduced in these cells, suggesting that the N551K-R1398H variant might induce LRRK2 protein instability in these model systems [36]. However, our findings do not reveal alterations in LRRK2 protein steady-state levels for the N551K-R1398H variant in human cells, suggesting that impaired protein stability is not a consistent phenotype of this LRRK2 variant. Our data and those of others indicate that LRRK2 GTP-binding, kinase activity, and Rab phosphorylation status do not fully explain the reduced PD risk in carriers of LRRK2 PD-protective variants. Further studies are warranted to determine whether the N551K, R1398H, or N551K-R1398H LRRK2 variants provide protection against neurodegeneration induced by familial *LRRK2* mutations *in vivo*, and if so, by what mechanism. Interestingly, replacement of R1398 with a leucine (R1398L) has been shown to increase GTP hydrolysis, reduce Rab10 phosphorylation induced by G2019S LRRK2 in cells, and attenuate dopaminergic neuronal loss induced by the adenoviral (Ad5)-mediated expression of G2019S LRRK2 in the rat nigrostriatal pathway [30, 47]. It remains to be determined how or whether R1398H, N551K, or their combination, can reduce PD risk, and at present their effects in cells and neurons tend to be subtle and varied.

Collectively, our study suggests that LRRK2-mediated Rab phosphorylation is a relevant indicator of the genetic risk associated with LRRK2 PD-coding variants in cells. The reduction of LRRK2 kinase activity as a therapeutic approach to slow disease progression in PD subjects harboring *LRRK2* PD risk variants is supported by the current study. Future studies are necessary to further elucidate the mechanisms by which LRRK2 PD-risk and PD-protective variants influence neuronal integrity and neurodegeneration in cultured neurons and animal models.

## Materials and Methods

### Animals

Time-pregnant female Sprague-Dawley outbred rats were obtained from Taconic Biosciences and P0-P1 rats were used to prepare post-natal primary cortical neuronal cultures as previously described [34]. Rodents were maintained in a pathogen-free barrier facility and provided with food and water *ad libitum* and exposed to a 12 h light/dark cycle. Animals were treated in strict accordance with the NIH Guide for the Care and Use of Laboratory Animals. All animal experiments were approved by the Van Andel Institute Institutional Animal Care and Use Committee (IACUC).

### Expression plasmids

Mammalian expression plasmids containing 3xFLAG-tagged full-length human LRRK2 variants (WT, R1441C, Y1699C, and G2019S) have previously been described [48]. LRRK2 missense mutations (A419V, N551K, R1398H, N551K-R1398H, R1628P, M1646T, S1647T, and G2385R) were introduced into 3xFLAG-LRRK2-WT by site-directed mutagenesis using the Stratagene QuickChange II XL kit (Agilent Technologies, La Jolla, CA, USA). LRRK2 double mutants (G2019S + common coding variant) were generated by introducing the G2019S mutation into the corresponding 3xFLAG-LRRK2 common coding variant by site-directed mutagenesis using the Stratagene QuickChange II XL kit. All expression plasmids were verified by DNA sequencing. A GFP-tagged Rab10 mammalian expression plasmid was previously described [49]. A list of expression plasmids used or generated are shown below.

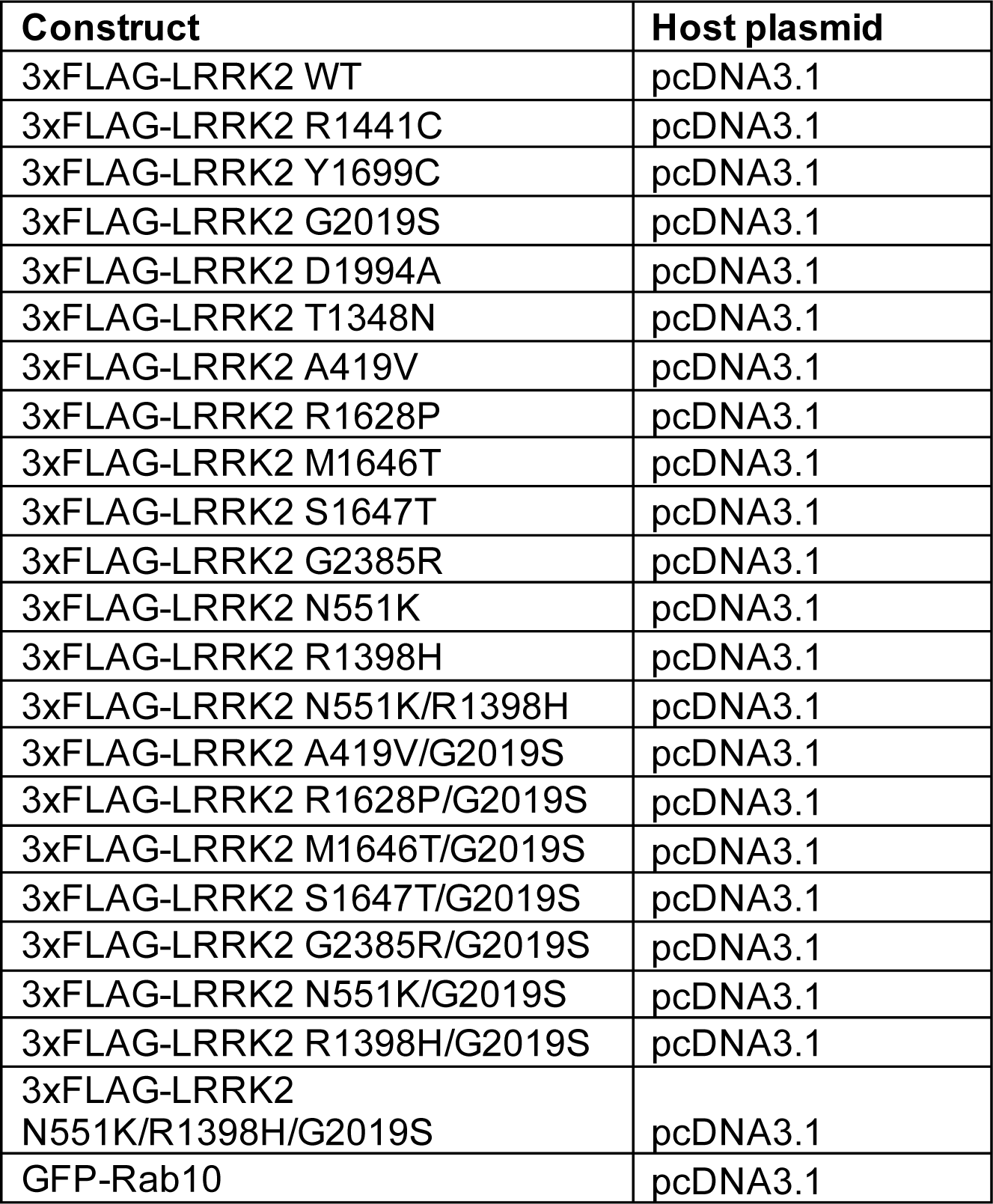

### Antibodies

For Western blot and immunocytochemical analysis, the following primary antibodies were used: mouse monoclonal anti-GFP (clones 7.1 and 13.1, Sigma), mouse monoclonal anti-FLAG and anti-FLAG-HRP (clone M2, Sigma), rabbit monoclonal anti-pSer910-LRRK2 (clone UDD1 15(3), Abcam), rabbit monoclonal anti-pSer935-LRRK2 (clone UDD2 10(12), Abcam), rabbit monoclonal anti-pSer1292-LRRK2 (clone MJFR-19-7-8, Abcam), rabbit monoclonal anti-phospho-Thr73-Rab10 (MJF-R21 or MJF-R21-22-5, Abcam), rabbit monoclonal anti-Rab10 (D36C4, Cell Signaling Technology), mouse monoclonal anti-β-tubulin (clone TUB 2.1, Sigma), mouse monoclonal anti-GAPDH (clone 1E6D9, Proteintech).

### Reagents

The LRRK2 kinase inhibitor, MLi-2 (Merck), was dissolved in DMSO and 1 mM stock solutions were stored at -80°C.

### Primary neuronal culture

Primary cortical neurons were plated in 35-mm dishes and maintained as described [50]. Sprague-Dawley pups were euthanized at P0-P1 and the cerebral cortices were stereoscopically isolated from whole brains. Brain tissues were dissociated in media containing papain (20 U/ml, Sigma). The cells were grown in 35-mm dishes on glass coverslips pre-coated with mouse laminin (33 mg/ml, ThermoFisher Scientific) and poly-D-lysine (20 ng/ml, Sigma) in media consisting of Neurobasal (ThermoFisher Scientific), B27 supplement (2% w/v, ThermoFisher Scientific), L-glutamine (500 µM, ThermoFisher Scientific) and penicillin/streptomycin (100 U/ml, ThermoFisher Scientific). At days-*in-vitro* 3 (DIV3), cells were treated with cytosine β-D-arabinofuranoside (AraC, 10 mM, Sigma) to inhibit glial cell division.

### Cell culture

Human SH-SY5Y and HEK-293T cells (ATCC) were maintained as described previously [50]. Cells were maintained in Dulbecco’s modified Eagle’s media (DMEM, Gibco) supplemented with 10% fetal bovine serum (ThermoFisher Scientific) and 1X penicillin/streptomycin (ThermoFisher Scientific) at 37°C in a 5% CO_2_ atmosphere. Cells were routinely passaged every 72 hours and maintained in culture up to the 30^th^ passage.

### Cell transfection

For biochemical analysis, HEK-293T cells at 2.5 x 10^5^ cells/condition were plated in 35-mm dishes 24 h before transfection. Transient transfection was performed with plasmid DNAs using XtremeGene HP DNA Transfection reagent (Roche) according to the manufacturer’s instructions. For LRRK2 steady-state levels, GTP-binding, and cellular and autophosphorylation assays, the cells were transfected using 2 µg of plasmids expressing full-length FLAG-tagged LRRK2 variants. At 48 h post-transfection, cells were washed twice in cold TBS (50 mM Tris pH 7.5, 150 mM NaCl) and harvested in 200 µl lysis buffer (50 mM Tris pH 7.5, 150 mM NaCl, 1% Triton-X100, 1X Complete Protease Inhibitor Cocktail [Roche], Phosphatase Inhibitor Cocktail Set 2 and 3 [Sigma]). For assessing Rab10 phosphorylation at Thr73, cells were co-transfected with 2 µg of each FLAG-tagged LRRK2 plasmid and 0.5 µg of GFP-tagged Rab10 plasmid. At 48 h post-transfection, cells were washed twice in cold TBS (50 mM Tris pH 7.5, 150 mM NaCl) and harvested in 200 µl lysis buffer (50 mM Tris pH 7.5, 150 mM NaCl, 1% Triton-X100, 1X Complete Protease Inhibitor Cocktail [Roche], Phosphatase Inhibitor Cocktail Set 1 [Sigma], 0.2 mM sodium orthovanadate, 10 mM sodium fluoride, 2 mM 2-glycerophosphate, 2 mM sodium pyrophosphate). Cell lysates were rotated at 4°C for 30 min and centrifuged at 21,000*g* for 15 min at 4°C to prepare supernatants. Protein concentration was determined using BCA assays (Pierce Biotech).

For immunocytochemistry, HEK-293T cells were plated on glass cover slips in 35-mm dishes at a density of 2.5 x 10^5^ cells/dish and transfected with FLAG-tagged LRRK2 variants as described above. At 48 h post-transfection, cells were fixed in 4% paraformaldehyde (PFA) and processed for immunofluorescence labeling. Primary cortical neuronal cultures at DIV 3 were transfected with FLAG-LRRK2 (4 µg total DNA per 35-mm dish) using Lipofectamine 2000 reagent (Invitrogen) according to the manufacturer’s recommendations. At DIV 6, cultures were fixed with 4% paraformaldehyde and processed for immunofluorescence labeling.

### Western blot analysis

Equivalent protein samples were mixed with 2X Laemmli sample buffer containing 5% 2-mercaptoethanol and resolved by SDS-PAGE using 7.5% Tris-Glycine (for FLAG-LRRK2) or 10% Tris-Glycine (for GFP-Rab10) gels followed by electrophoretic transfer onto nitrocellulose membranes (0.2 µm, GE Life Sciences). Membranes were blocked in TBS and 0.1% Tween-20 with 5% non-fat milk (Bio-Rad) for 1 h at room temperature, and incubated with primary antibodies overnight at 4°C. For assessing cellular phosphorylation or autophosphorylation of LRRK2 (pSer935, pSer910, and pSer1292) and Rab10 (pThr73), equivalent protein samples were loaded into separate gels of total protein (FLAG-LRRK2 or GFP-Rab10) to avoid sequential stripping. Secondary antibodies conjugated to HRP were detected using enhanced chemiluminescence (ECL, GE Life Sciences) with images captured on a luminescent image analyzer (LAS-3000, Fujifilm). Protein bands were quantified by densitometry using Image Studio™ Lite, v4.0 software.

### Immunocytochemistry and confocal microscopy

For immunocytochemistry, coverslips were incubated with primary antibodies diluted at 1:200 in PBS and 0.3% Triton-X100 buffer supplemented with 5% horse serum, 5% goat serum, and 30 mg/ml BSA overnight at 4°C in a humid chamber. Coverslips were washed three times in PBS and incubated with secondary antibodies diluted at 1:500 in PBS for 1 h at room temperature. Coverslips were again washed three times in PBS and were mounted on glass slides (Fisherbrand Premium) using ProLong™ Gold antifade mountant with DAPI (ThermoFisher Scientific). Fluorescent images were acquired using a Nikon A1 Plus inverted confocal microscope using NIS-Element software (Nikon). Representative images were taken from a single *z*-plane at a thickness of 0.1 mm.

### Lysosomal stress assays

Nigericin (InvivoGen), monensin (M5273, Sigma), Leu-Leu methyl ester hydrobromide (LLOMe, L7393, Sigma), and chloroquine (C6628, Sigma) were used to induce lysosomal stress and stock solutions were prepared according to the manufacturer’s instructions. For lysosomal stress assays, cells were co-transfected with FLAG-LRRK2 and GFP-Rab10 plasmid as described above. At 48 h post-transfection, cells were treated with nigericin (2 µM), monensin (10 µM), LLOMe (1 mM), and chloroquine (50 µM) by media change and harvested at the indicated time for cell lysis as described above.

### GTP-binding assay

SH-SY5Y cell lysates were collected as above. Equivalent amounts of protein sample were incubated with 30 µl of guanosine 5′-triphosphate-agarose (Sigma-Aldrich) in PBST buffer (1X PBS, 1% Triton X100) to a final volume of 1 ml and subjected to GTP-binding by rotating for 2 h at 4°C. Agarose beads were washed 3 times with PBST and 2 times with 1X PBS. GTP-bound fractions were eluted in Laemmli buffer containing 5% 2-mercaptoethanol and heated at 70°C for 10 min. GTP-bound fractions and input lysates (1% total lysate) were resolved by SDS-PAGE (7.5% Tris-Glycine gels) and subjected to Western blot analysis with anti-FLAG-HRP antibody (Sigma).

### Neurite outgrowth assay

Neurite outgrowth assays were performed using methods as previously described [34]. Primary cortical cultures at DIV 3 were co-transfected with FLAG-LRRK2 and GFP plasmids at a 10:1 molar ratio (5 µg total DNA per 35-mm dish) using Lipofectamine 2000 reagent (Invitrogen) according to the manufacturer’s recommendations. At DIV 6, cultures were fixed with 4% paraformaldehyde and processed for immunocytochemistry with primary antibodies (mouse anti-FLAG-M2 antibody [Sigma] and rabbit anti-βIII-tubulin [Sigma]) and secondary antibodies (anti-mouse IgG-AlexaFluor-633 and anti-rabbit IgG-AlexaFluor-594 [Invitrogen]). Fluorescent images were acquired using an Olympus fluorescence digital microscope (BX61) at 20X for neurite length measurements. Only neurons that had extended neurites were measured whereas neurons without processes were excluded from the analysis. GFP-positive neurites were assigned as axons (longest neurite) or dendrites (all neurites minus the longest neurite). Neurites were traced using NIH ImageJ software.

### Statistical Analysis

All statistical analyses were performed with GraphPad Prism software. One-way ANOVA with appropriate *post-hoc* analysis was used for multiple comparisons, whereas unpaired, two-tailed Student’s *t*-test was used for comparing two groups, as indicated. All data were plotted as mean ± SEM and were considered significant when *P*<0.05.

## Acknowledgements

This study was supported by a grant from the National Institutes of Health (NIH) R01NS120489 to D.J.M., and by financial support from the Van Andel Institute.

## Acknowledgements

We thank the VAI Optical Imaging Core (RRID:SCR_021968) for technical assistance.

## Supplementary Figure Legends

**Figure S1. Impact of coding risk variants on LRRK2 subcellular localization in human cells.** HEK-293T cells transiently expressing FLAG-tagged LRRK2 risk variants or familial mutations were fixed at 48 h post-transfection and subjected to immunofluorescence. Representative confocal images taken from a single z-plane at 0.1 mm thickness showing LRRK2 variants (FLAG, green), endogenous pThr73-Rab10 (red), and nuclei (DAPI, blue). Note, LRRK2 variants are mostly cytoplasmic in cells and do not form skein-like filamentous structures. Scale bar: 10 µm.

**Figure S2. Impact of coding risk variants on LRRK2 subcellular localization in cortical neurons.** Rat primary cortical neurons were transfected at DIV 3 with FLAG-tagged LRRK2 risk variants or familial mutations and fixed at DIV 6 for immunofluorescence analysis. Representative confocal images taken from a single z-plane at 0.1 mm thickness showing LRRK2 variants (FLAG, green), endogenous pThr73-Rab10 (red), and nuclei (DAPI, blue). Note, LRRK2 variants are mostly cytoplasmic in neurons and do not form skein-like filamentous structures. Scale bar: 10 µm.

**Figure S3. Effects of pharmacological kinase inhibition on LRRK2 subcellular localization in cortical neurons.** Rat primary cortical neurons were transfected at DIV 3 with FLAG-tagged LRRK2 risk variants or familial mutations and treated with the LRRK2 kinase inhibitor MLi-2 (100 nM) or PBS (control) for 3 hours before fixation at DIV 6. Representative confocal images taken from a single z-plane at 0.1 mm thickness showing LRRK2 variants (FLAG, green), pSer935-LRRK2 (red), and nuclei (DAPI, blue). Note, the complete absence of pSer935-LRRK2 signal following MLi-2 treatment, thereby confirming LRRK2 kinase inhibition. LRRK2 variants are mostly cytoplasmic and do not form skein-like filamentous structures following kinase inhibition. Scale bar: 10 µm.

**Figure S4. PD risk and protective LRRK2 coding variants support increased Rab10 phosphorylation induced by lysosomal stressors.** Western blot analysis of GFP-Rab10 phosphorylation at Thr73 and LRRK2 phosphorylation at Ser935 and Ser1292 induced by FLAG-tagged LRRK2 risk variants or familial/synthetic mutations in SH-SY5Y cells treated with the lysosomal stressors (A) 10 µM monensin for 2 h or (B) 1 mM LLOMe for 1 h relative to PBS (control). (A-B) Bars represent mean ± SEM fold induction of pThr73-Rab10, pSer1292-LRRK2, or pSer935-LRRK2 for each LRRK2 risk variant by each lysosomal agent relative to control treatment (*n* = 3-5 independent transfections). *P*>0.05 compared with WT LRRK2 by multiple Student’s *t*-test.

## Notes

### Competing Interest Statement

The authors have declared no competing interest.

