## Supplementary figures and images for "Parkinson’s disease-associated LRRK2 risk variant, G2385R, enhances Rab substrate phosphorylation and impairs neuronal integrity"

### Figure S1

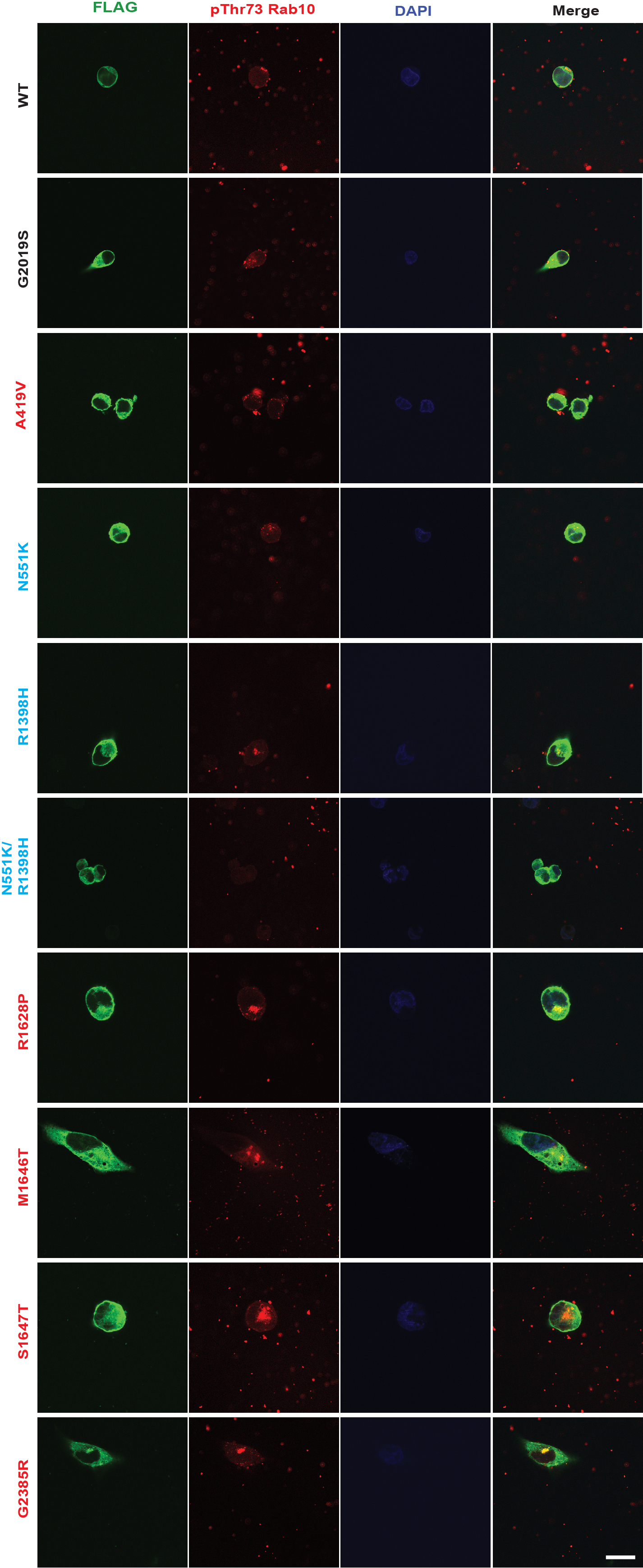

### Figure S2

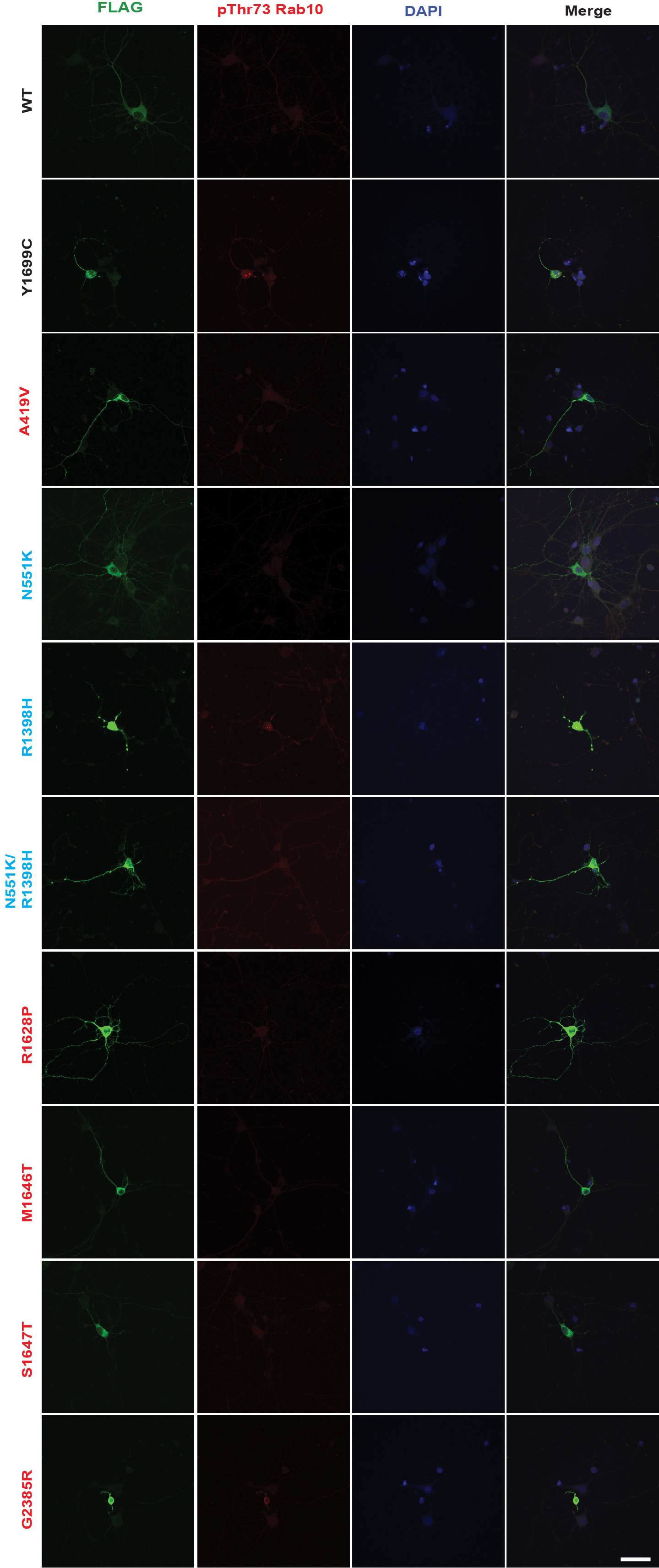

### Figure S3

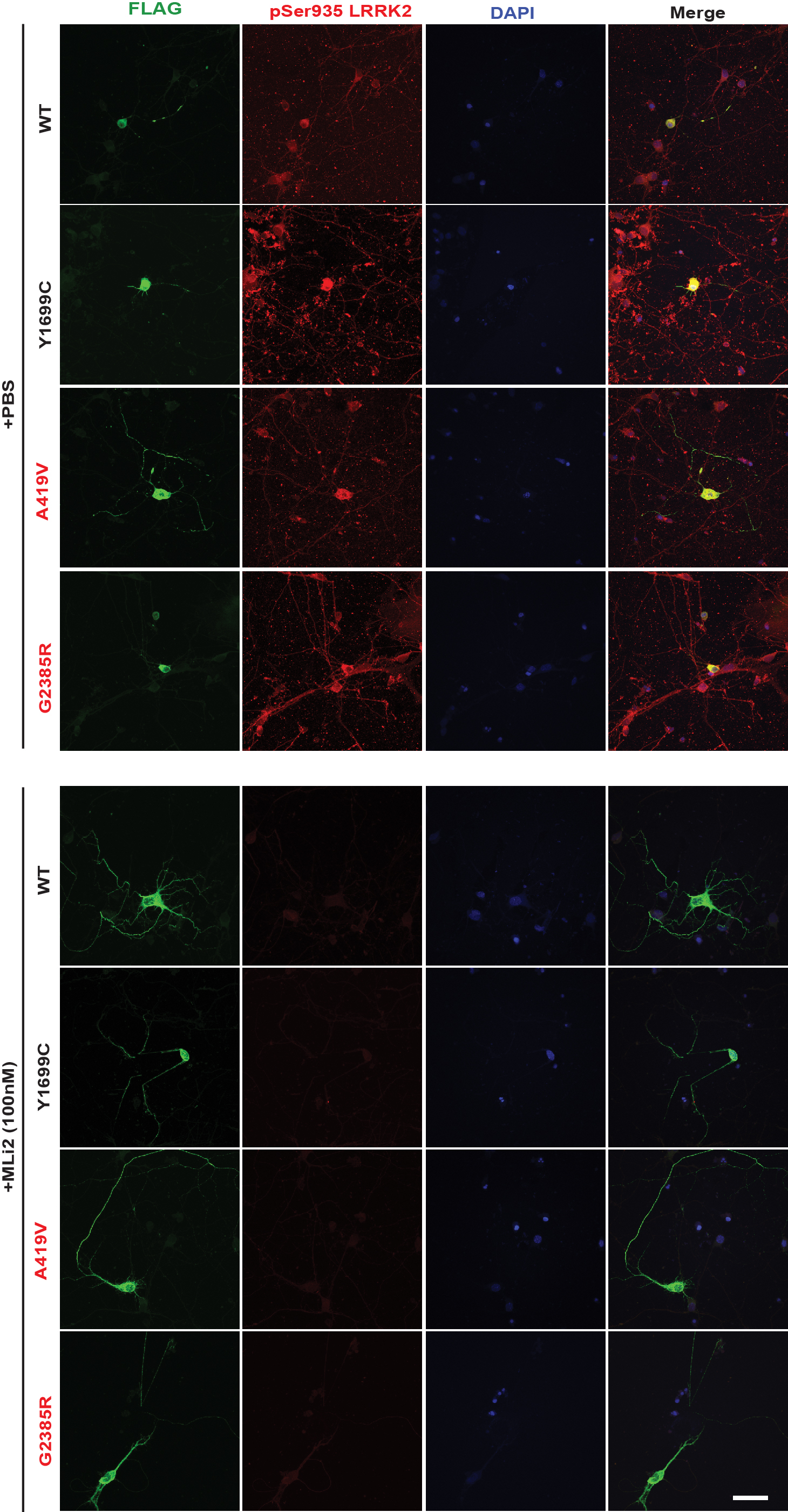

### Figure S4

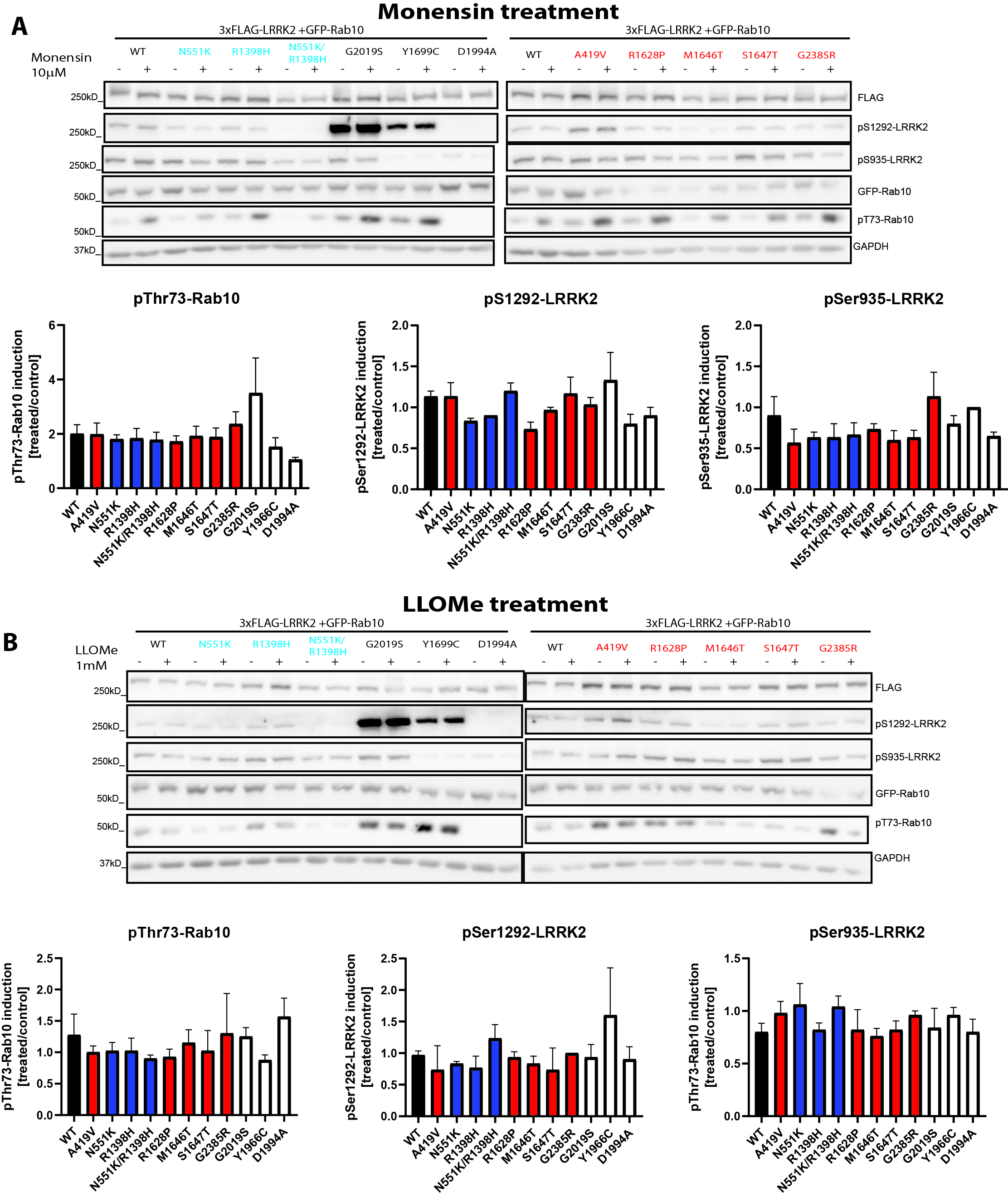
